# Self-organization of a functional human organizer by combined WNT and NODAL signalling

**DOI:** 10.1101/234633

**Authors:** I. Martyn, T.Y. Kanno, A. Ruzo, E.D. Siggia, A.H. Brivanlou

## Abstract

In amniotes, the development of the primitive streak (PS) and its accompanying “organizer” define the first stages of gastrulation. Despite detailed characterization in model organisms, the analogous human structures remain a mystery. We have previously shown that when stimulated with BMP4, micropatterned colonies of human embryonic stem cells (hESCs) self-organize to generate early embryonic germ layers^1^. Here we show that in the same type of colonies WNT signalling is sufficient to induce a PS, and WNT with ACTIVIN is sufficient to induce an organizer, as characterized by embryo-like sharp boundary formation, epithelial-to-mesenchymal transition (EMT) markers, and expression of the organizer specific transcription factor GSC. Moreover, when grafted into chick embryos, WNT and ACTIVIN treated human cells induce and contribute autonomously to a secondary axis while inducing neural fate in the host. This fulfills the most stringent functional criteria for an organizer, and its discovery represents a major milestone in human embryology.

---

The pioneering experiments that Spemann and Mangold published in 1924 demonstrated that a small group of cells located on the dorsal side of the early amphibian embryo have the ability to induce and “organize” a complete secondary axis when transplanted to the ventral side of another embryo^2^. Thus the “organizer” concept was born, and later discovery of embryonic tissue with similar organizer activity in fish, birds, and rodents^3,4,5,6^ demonstrated that this early embryonic activity was evolutionarily conserved. Cells in the organizer of all species so far studied display the same behavior during axis induction: (i) they contribute autonomously to axial and paraxial mesoderm, including head process and notochord; and (ii) induce neural fate non-autonomously in their neighbors. The organizer in primates, and especially humans, has so far not been defined.

Due to the ethical limitations of working with early human embryos, the only way to search for the human organizer is via human embryonic stem cells (hESCs). We have previously shown that when grown on geometrically confined disks hESCs respond to BMP4 by differentiating and self-organizing into concentric rings of embryonic germ layers: with ectoderm in the center, extra-embryonic tissue at the edge, and mesoderm and endoderm in between^1^. These embryo-like “gastruloids” are robust and amenable to analysis with subcellular resolution.

During mouse gastrulation BMP4 signalling activates the WNT pathway which in turn activates the ACTIVIN/NODAL pathway (BMP4→WNT→ACTIVIN/NODAL, Fig. 1A), both at the transcriptional and signalling level^7^. Since it has been shown in mouse and other vertebrates that these three pathways are the most critical for organizer formation^8,9,10,11^, we first asked if this hierarchy was conserved in human gastruloids. Using RNA-Seq, we found that out of all the 19 *WNT* ligands present in the human genome, *WNT3* is the only one that is significantly and immediately induced upon BMP4 presentation (Fig. 1B). qPCR analysis shows that activation of WNT signalling directly induces *NODAL* expression (Fig. 1C). Further qPCR analysis showed that NODAL induction was reduced when the NODAL inhibitor SB was present, and taken together with the observation that ACTIVIN induces *NODAL* expression, suggests the presence of a NODAL feedback loop, as also noted in the mouse^12^. Additionally, no direct *BMP4* induction by either WNT or NODAL signalling was observed (Supp. Fig. 1B). Thus the transcriptional hierarchy of BMP→WNT→NODAL is evolutionarily conserved in hESCs.

**Figure 1.**
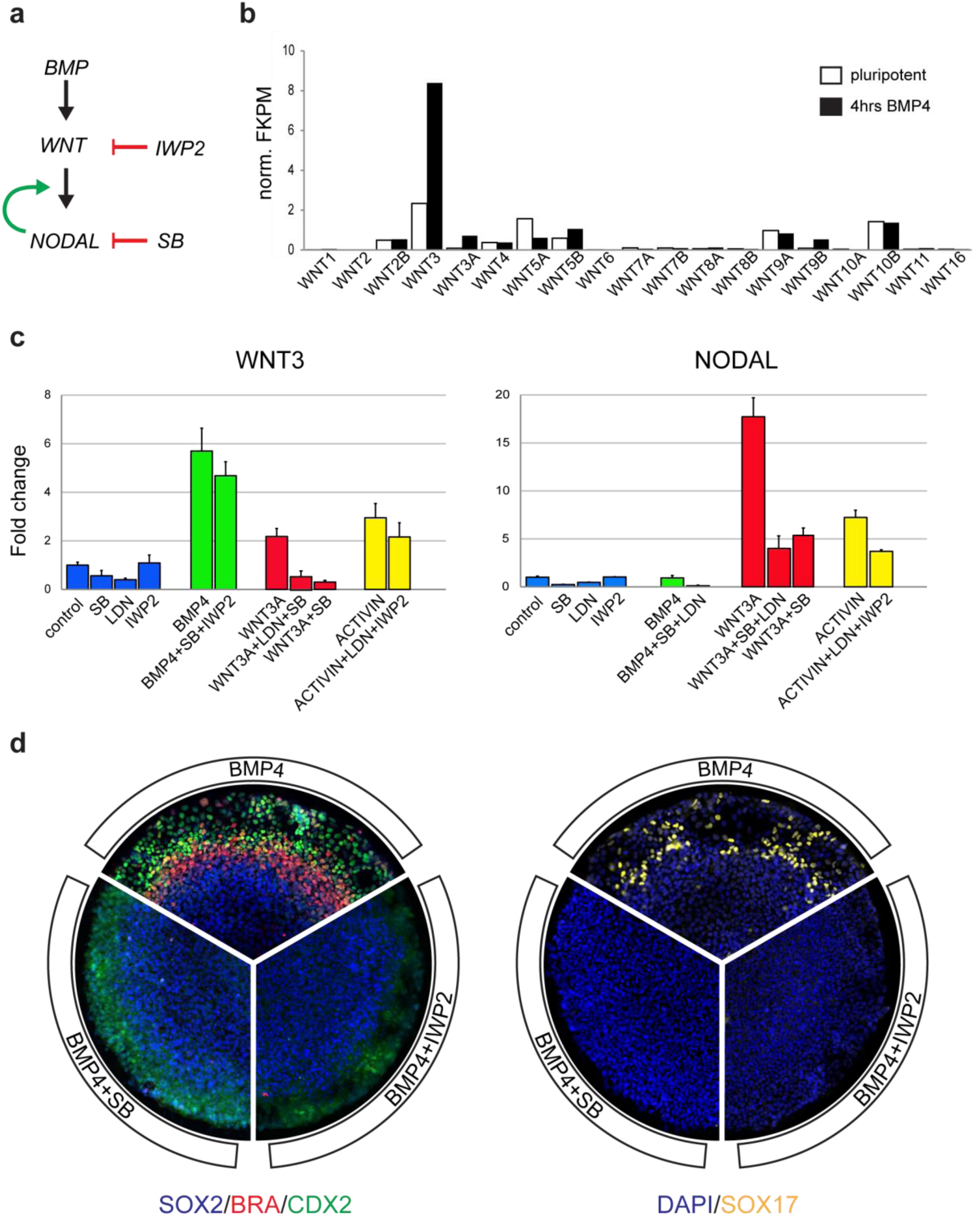
hESCs obey BMP->Wnt->Nodal PS signalling initiation hierarchy. (a) Model of proposed PS signalling initiation hierarchy in hESCs, along with indication at which step the inhibitors SB and IWP2 act. Like in mouse, BMP acts on WNT, and then WNT acts on NODAL. There is also positive feedback between WNT and NODAL. (b) RNA-seq expression of all known WNT ligands in pluripotency and after 4hrs of BMP4 in hESCs on 500μm diameter micropatterns. The results show that overall WNT transcription is low in pluripotency and that WNT3 is the only strong and direct WNT ligand target of BMP4 stimulation. (c) qPCR of WNT3 and NODAL in small colonies of hESCs after 4hrs stimulation with each condition shown on x-axis. The data is consistent with the predictions of the model. (d) Pie sections are of representative 1000μm diameter micropatterned colonies stimulated with BMP4, BMP4+IWP2, or BMP4+SB and fixed and stained for germ layer molecular markers after 48hrs. All micropattern experiments were performed on at least 3 separate occasions, and unless mentioned otherwise, all other micropatterns are 1000μm in diameter. Staining is quantified in Supp. Fig. 1C.

To test if the hierarchy of signalling activity was also conserved, we challenged BMP4 self-organizing activity with two small inhibitors: SB and the WNT inhibitor IWP2. When presented alone neither of the inhibitors had any effect, but challenging BMP4 activity by either IWP2 or SB led to a loss of mesoderm (BRA) and endoderm (SOX17; Fig. 1D and quantified in Supp. Fig. 1C). Thus both WNT and ACTIVIN/NODAL signaling are necessary for mesendodermal induction and patterning downstream of BMP4.

To ask if WNT or ACTIVIN/NODAL signalling alone were sufficient to induce differentiation and self-organization, and if they could yield a population of cells with an organizer identity, hESC colonies were stimulated with either WNT3A or ACTIVIN. After 48 hours of treatment, WNT3A led to differentiation of the periphery into mesoderm (BRA) and endoderm (SOX17; Fig. 2A and quantified in Supp. Fig. 2A). The center cells maintained their pluripotent epiblast fate (SOX2 and NANOG) rather than differentiating into ectoderm (Fig. 2A and B). After 48 hours of ACTIVIN treatment, however, no cells showed any sign of differentiation or self-organization, and all maintained the same morphology and expression of the pluripotency markers (Fig. 2A and B).

**Figure 2.**
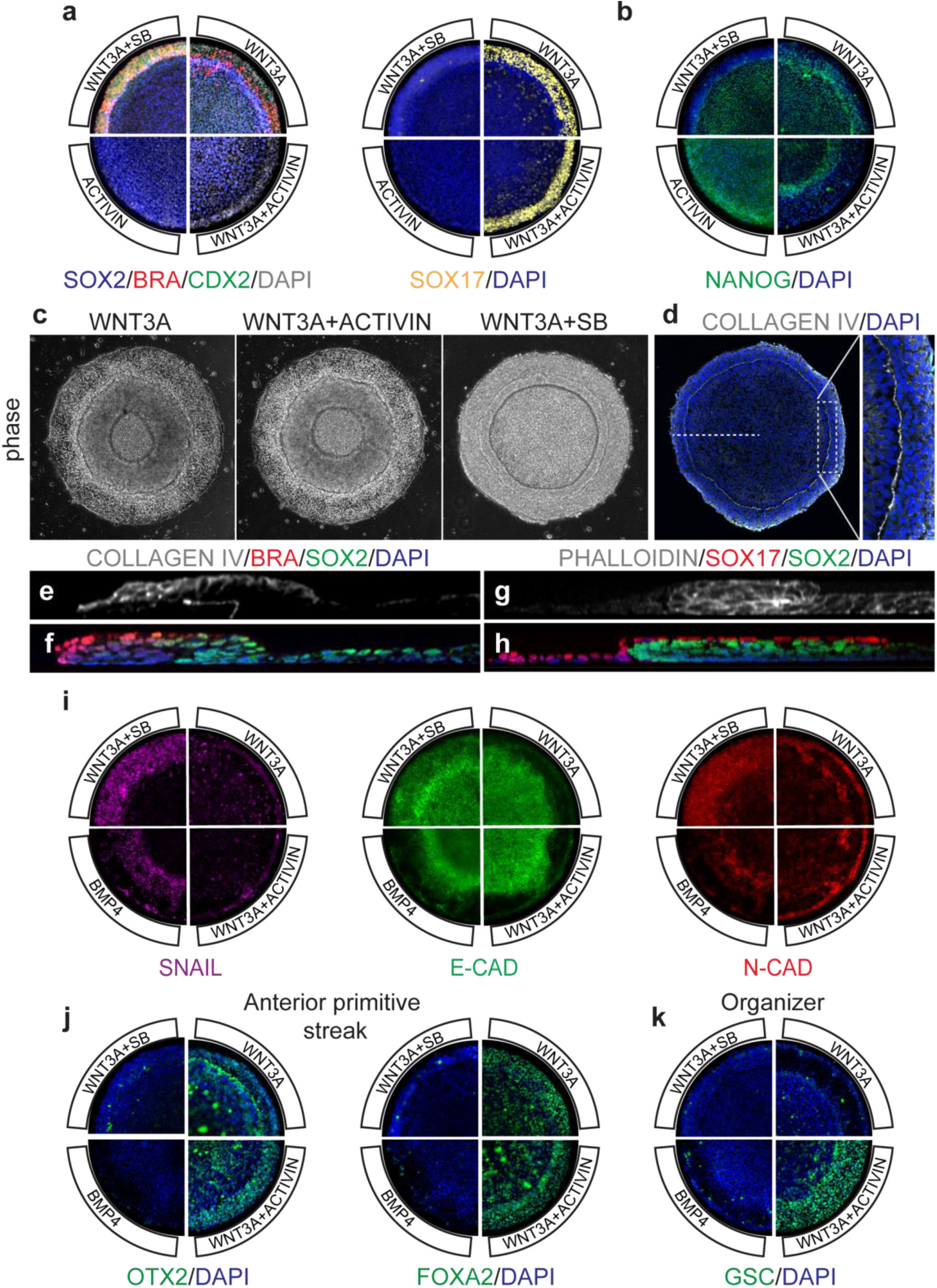
WNT is necessary and sufficient to induce PS markers and morphology. (a) Micropatterned colonies stimulated with WNT3A, WNT3A+ACTIVIN, WNT3A+SB, or ACTIVIN and fixed and stained for germ layer molecular markers after 48hrs. Staining is quantified in Supp. Fig. 2A. (b) Micropatterned colonies stimulated with WNT3A, WNT3A+ACTIVIN, WNT3A+SB, or ACTIVIN and fixed and stained for pluripotency marker NANOG after 48hrs. (c) Sharp boundaries are visible even in phase contrast of WNT3A, WNT3A+SB, and WNT3A+ACTIVIN micropatterns at 52hrs. (d-f) Micropatterns stained with DAPI (blue; d and f), COLLAGEN IV (white; d and e), BRA (red; f), and/or SOX2 (green; f) after 52hrs of WNT3A+SB. Note how COLLAGEN IV traces a sharp continuous circle between the periphery and the interior of colony and forms a dividing line between the epiblast-like SOX2+ interior and differentiating BRA+ primitive streak exterior. In addition, our EMT data shows that the BRA- and SOX2-cells express SNAIL. (g) and (h) Micropatterns stained with phalloidin (white; g) or DAPI (blue; h), SOX17 (red; h), and SOX2 (green; h) after 52hrs of WNT3A. Cross section shows an inner epithelial SOX2+ epiblast region folded back in on itself. Outside and above the fold is the mesenchymal SOX17 region. Such geometry seems to permit the mesenchymal SOX17 cells to migrate along the epiblast-like region in basal-to-basal contact. Corroborating this, no SOX17 cells were ever observed further inwards beyond the leading edge of the fold where the basal presenting surface ends. (i) Micropatterned colonies stimulated with BMP4, WNT3A, WNT3A+SB, or WNT3A+ACTIVIN and fixed and stained for EMT markers SNAIL, E-CAD, and N-CAD after 48hrs. Note that WNT3A and WNT3A+ACTIVIN stimulated colonies show less SNAIL at 48hrs because they started their EMT earlier, at 24hrs (see Supp. Fig. 2B for 12, 24, and 36hr timepoints). (j-k) Micropatterned colonies stimulated with WNT3A, WNT3A+ACTIVIN, WNT3A+SB, or BMP4 and fixed and stained after 48hrs for marker set characteristic of anterior PS (OTX2 and FOXA2; j) or organizer specific transcription factor GSC (k). Stainings are quantified in Supp. Fig. 3A.

As it is unlikely that ACTIVIN/NODAL has no effect during human gastrulation, we presented WNT3A in two combinations that represent the opposite extremes of a ACTIVIN/NODAL gradient: WNT3A+ACTIVIN and WNT3A+SB. In accordance with studies in model systems and human and mouse embryonic stem cells^13,14^, we find that ACTIVIN/NODAL signalling acts as a modifier of mesoderm and endoderm patterning, with all the cells on the periphery converting to endoderm (SOX17+) with no mesoderm (BRA-) in WNT3A+ACTIVIN, and all cells converting to mesoderm (BRA+) with no endoderm (SOX17-) in WNT3A+SB (Fig. 2A and Supp. Fig. 2A). As with WNT3A treatment alone, both sets of colonies had sharp morphological boundaries that became much more pronounced after 48 hours (Fig. 2C). The expression profile of COLLAGEN IV, which in mouse demarcates the basement membrane between the epiblast and the visceral endoderm and which breaks down as the PS progresses^15^, and the 3D organization of cells around this boundary (Fig. 2D-H), was highly reminiscent of the morphological signature of a PS. Confirming the PS nature of these structures, we find evidence of an epithelial-to-mesenchymal transition (EMT) with SNAIL expression and an E-CADHERIN (E-CAD) to N-CADHERIN (N-CAD) switch where the mesendodermal fates later establish themselves (Fig. 2I and Supp. Fig. 2B). Taken together, we find that WNT signalling is necessary and sufficient to induce PS, and that ACTIVIN/NODAL signalling acts as a modifier that controls timing of EMT and patterning of mesoderm versus endoderm.

Having identified PS in our human gastruloids, we next asked whether an organizer subpopulation is also present. In mouse, the organizer is located in the anterior PS in what is thought to be the highest region of NODAL signalling^16^. We find that both WNT and WNT3A+ACTIVIN treatment results in co-expression of OTX2 and FOXA2 in the same micropattern region that expresses SOX17, this set of markers together being characteristic of anterior PS in mouse. However, as in the mouse, only the condition with the highest NODAL signalling, i.e. WNT3A+ACTIVIN, results in the expression of the organizer-specific marker GSC (Fig. 2J-K, and quantified in Supp. Fig. 3C). WNT3A+ACTIVIN also leads to the highest expression of key secreted inhibitors known to be produced by the organizer and its deriviatives^17^, such as CHORDIN, DKK1, CER1, LEFTY1, and LEFTY2, and it also leads to the highest expression of NODAL, which at later stages in mouse is also specific to the organizer (Supp. Fig. 4D).

The induction of characteristic organizer markers, the emergence of a sharp COLLAGEN IV based morphological boundary dividing the PS and epiblast regions, and the induction of EMT all provide evidence in support of the induction of a human organizer in an early primitive streak by WNT3A+ACTIVIN treatment. However, as originally defined by the classic amphibian experiments, an organizer is determined functionally as a group of cells that can induce a secondary axis when grafted ectopically into host embryos^2^. In this context, the grafted cells should contribute directly to the ectopic axis (autonomously), and induce neural tissue in the cells of the host (non-autonomously). In order to test for the most stringent and functional definition of an organizer, we used an *ex ovo* cross-species transplantation strategy based on previous mammalian organizer studies^18,19^, grafting fluorescent reporter hESC micropatterns treated for 24 or 48 hours with WNT3A+ACTIVIN into the marginal zone of Early Chick (EC) culture embryos^20^ (stage HH 2 to 3+). We used 500μm diameter rather than 1000μm diameter micropatterns as these gave a purer population of GSC+ cells, and we grafted at 24 hours post-treatment as well as at 48 hours post-treatment as 24 hours is when GSC first becomes apparent and is also co-expressed with BRA (Supp. Fig. 4A-C). For the reporter line, we used the CRISPR-Cas9 generated RUES2-GLR (Germ Layer Reporter) cell line, as it encodes 3 separate fluorescent markers at the endogenous locations for each of the 3 germ layers (see Supp. Fig. 5 and 6).

We found that RUES2-GLR grafts survived, mingled with host cells, and induced and contributed to a secondary axis that became obvious between 24-48 hours (Fig. 3B-L). Both the live cell reporter and a human-specific nuclear antigen revealed that the human cells directly contributed to the ectopic axis autonomously and continued differentiating in their new location, contributing both BRA and SOX17 cells (Fig. 3H, L-M). This mirrors previous observations in mouse-to-mouse organizer grafting experiments^5^. Confocal cross-sectioning of these secondary axes often revealed self-organizing features directly resembling those found in the early chick and mouse embryo, for example correct layering of germ layers, and central elongated notochord-like structures (Fig. 3 N-R and Supp. Video 1). Analysis of molecular markers also established that the human cells induced neural tissue in the chick non-autonomously: SOX2 and SOX3 were ectopically induced in chick cells that surrounded the human cells (Fig. 3E-G, I-K, S). Additional *in situs* and antibody staining for HOXB1, GBX2, and OTX2 established that the neural tissue was predominantly posterior in nature (Fig. 3 T-V). Since in the mouse the early-gastrula-organizer (EGO) and late-streak node also does not induce anterior neural structures when grafted to another mouse embryo, this result suggests that our human organizer is closer to these organizer stages than to the mouse mid-gastrula-organizer (MGO) ^6,17^. As controls, RUES2-GLR grafts treated instead with WNT3A, WNT3A+SB, BMP4, or blank media showed less overall survival and never induced chick neural markers (Table 1 and Supp. Fig. 5A). Taken together with the morphological, cellular, and molecular evidence described above, this functional test in an embryonic environment provides the most stringent evidence for the induction of a human organizer. It also highlights that the organizer itself can be obtained *in vitro* by self-organization of hESCs in response to WNT+ACTIVIN treatment in a confined micropattern geometry.

**Figure 3.**
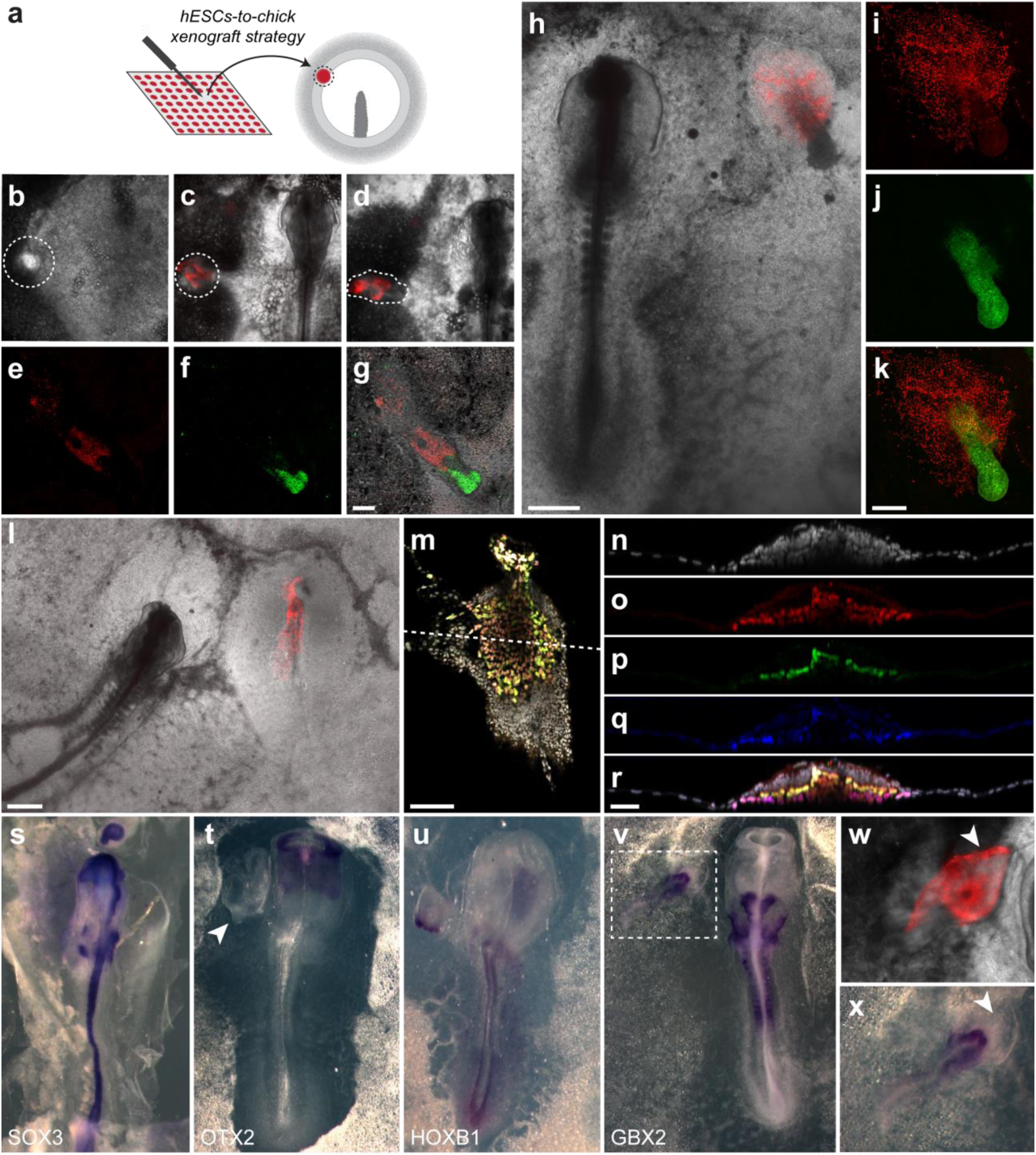
Human organizer induces secondary axis in chick embryo. (a) Schematic showing strategy for grafting 500μm diameter WNT3A+ACTIVIN induced hESC micropatterns into the marginal zone of HH stage 2-3+ chick embryos, ∼90° away from site of host PS initiation. (b-g) Secondary axis induced by 24hr stimulated RUES2-GLR colony into HH stage 3 chick; (b-d) SOX17-tdTomato (red) live marker at 0, 24, and 38hrs post-graft (unfortunately this was the only marker that could be followed live as the others were obscured by the agar mount and chick background auto-fluorescence); (f-g) SOX2 (green) is prominent in the tip of the secondary axis 48 hrs post-graft, and does not overlap with the hESCs (e and g, red, Human Nuclear Antigen). Scale bar is 200μm. (h-k) Another example of a 24hr stimulated hESC micropattern inducing a secondary axis in a chick host, 27hrs post-graft; (h) live image of SOX17-tdTomato hESC cells (red); (i and k) fixed stains for Human Nuclear Antigen (HNA, red) and SOX2 (j and k, green). Scale bars are 500μm (h) and 200μm (i-k). (l-r) Example of secondary axis induction from a 24 hr stimulated hESC micropattern with more complete self-organizing structures, 27hrs post-graft; (l) live image of SOX17-tdTomato hESC cells (red); (m) confocal slice of secondary axis for DAPI (grey), HNA (red), and BRA (green); (n-r) confocal cross-section of indicated region in (m), with the same channels plus SOX17 (blue). Note in the merged image (r) how the secondary axis is layered, with epiblast chick cells on top of a layer of human BRA cells which in turn are on top of a layer of human SOX17 cells, exactly how the epiblast, mesoderm, and endoderm layers would arrange themselves in a gastrulating mouse or chick embryo. Scale bars are 500μm (l), 100μm (m), and 50μm (n-r). (s) In situ for chicken *SOX3* shows expression in the host chick throughout the neural tube and head, as well as in the induced secondary axis. (t) *OTX2* is expressed in the host forebrain but is absent in the graft induced tissue (indicated by arrow). (u) *HOXB1* is expressed in the host and the graft induced secondary axis. (v) *GBX2* is expressed in the host and the graft induced secondary axis. (w-x) Zoom of region indicated in (v): (w) shows secondary axis and tdTomato-hESCs (red) after fixation; (x) shows *GBX2* expression after in situ. The arrow shows the location of the graft hESCs before and after.

**Table 1.** Induction of chick neural tissue by hESC micropatterns. Statistical analysis (χ^2^, 2×2contingency test, compared to the control condition): *P<0.05; **P<0.01; ***P<0.001; ****P<0.0001.

|  | **Survival** | **SOX2** | **SOX3** | **GBX2** | **HOXB1** | **OTX2** |
| --- | --- | --- | --- | --- | --- | --- |
| **control (blank)** | 4/15 | 0/15 | - | - | - | - |
| **WNT3A+ACTIVIN 24hrs** | 273/300**** | 10/19**** | - | 5/6 | 9/10 | 0/14 |
| **WNT3A+ACTIVIN 48hrs** | 37/40**** | 6/15**** | 6/8 | - | - | - |
| **WNT3A 48hrs** | 8/15 | 0/15 | - | - | - | - |
| **WNT3A+SB 48hrs** | 7/15 | 0/15 | - | - | - | - |
| **BMP4 48hrs** | 14/15*** | 0/15 | - | - | - | - |

Our ability to generate a functional human organizer closes the loop initiated by classical experimental embryologists working on amphibian systems, nearly 100 years ago, and demonstrates that the concept of the “organizer” is evolutionarily conserved from frogs to humans. Our chick experiments also define a novel *in vivo* platform to validate results obtained in an *in vitro* gastruloid platform, and may be generally applicable to test and explore other aspects of early human development.

## Acknowledgements

The authors are grateful to Irene Yan and Felipe Vieceli for materials and protocols, to J. Metzger for assistance with 3D image segmentation, and to members of the A.H.B. and E.D.S. laboratories for helpful discussions. This work was supported by grants R01 HD080699 and R01 GM101653.

## Author Contributions

Conceptualization and writing, I.M., A.H.B., and E.D.S.; Stem cell experiments performed by I.M., chick experiments devised by I.M. and performed by I.M. and T.Y.K.; Conception, generation, and validation of RUES2-GLR cell line by A.R.; All authors reviewed the manuscript.

## METHODS

### Cell Culture

hESCs (RUES2 cell line) were grown in HUESM medium conditioned by mouse embryonic fibroblasts (MEF-CM) and supplemented with 20ng/mL bFGF. Mycoplasma testing was carried out before beginning experiments and again at 2-month intervals. For maintenance, cells were grown on GelTrex (Invitrogen) coated tissue culture dishes (BD Biosciences, 1:40 dilution). The dishes were coated overnight at 4°C and then incubated at 37°C for at least 20 minutes before the cells were seeded on the surface. Cells were passaged using Gentle Cell Dissociation Reagent (Stem Cell Technologies 07174).

### Micropatterned Cell Culture

Micropatterned coverslips were made according to a new protocol devised in our lab that significantly reduced operating costs. First, 22×22mm #1 coverslips were spin-coated with a thin layer of PDMS (RTV615A Momentive) and left to set overnight. They were then coated with 5μg/ml laminin 521 (Biolamina) diluted in PBS with calcium and magnesium (PBS++) for 2 hours at 37°C. After 2 washes with PBS++, coverslips were placed under a positive feature UV Quartz Mask (Applied Image In) in a home-made UV oven. Laminin not protected by the features in the mask was burned off by 10 minutes of deep UV application (185nm wavelength). Coverslips were then removed, washed twice more with PBS++, and then left at 4°C overnight in 1% F127-Pluronic (Sigma) solution in PBS++. The now patterned coverslips were used within 1 week of fabrication. To seed cells onto a micropatterned coverslip, cells were dissociated from growth plates with StemPro Accutase (Life Technologies) for 7 minutes. Cells were washed once with growth media, washed again with PBS, and then re-suspended in growth media with 10μM ROCK-inhibitor Y-27632 (Abcam).

Coverslips were placed in 35mm tissue culture plastic dishes, and 1×10^6^ cells in 2mL of media were used for each coverslip. After 1 hour ROCK-inhibitor was removed and replaced with standard growth media supplemented with Pen Strep (Life Technologies). Cells were stimulated with the following ligands or small molecules 12hrs after seeding: 200ng/mL Wnt3a, 50ng/mL BMP4, 100ng/mL Activin-A, 10μM SB, or 2μM IWP2.

### hESC immunocytochemistry

Cells were fixed in 4% paraformaldehyde for 20 minutes at room temperature, washed twice with PBS, then blocked and permeabilized with 3% donkey serum and 0.1% Triton X-100 in PBS for 30 minutes, also at room temperature. Cells were incubated overnight with primary antibodies in this blocking buffer at 4°C. The next day they were washed 3 times with PBS+0.1% Tween-20 for 30 minutes each, then incubated with secondary donkey antibodies (Alexa 488, Alexa 555, Alexa 647) and DAPI for 30 minutes. Cells were then washed twice more with PBS, and then mounted on glass slides for imaging. Primary antibodies used are listed in Supplemental Table 1.

**Supplemental Table 1:** Antibody information

| Antigen | Antibody | Dilution |
| --- | --- | --- |
| BRACHYURY | R&D Systems AF-2085 | 1:300 |
| CDX2 | Abcam Ab-15258 | 1:50 |
| E-CADHERIN | Cell Signalling 3195 | 1:200 |
| GOOSECOID | R&D AF-4086 | 1:100 |
| NANOG | R&D Systems AF-1997 | 1:200 |
| N-CADHERIN | BioLegend 350802 | 1:200 |
| SNAIL | R&D Systems AF-3639 | 1:200 |
| SOX17 | R&D Systems AF-1924 | 1:200 |
| SOX2 | Cell Signalling 3579 | 1:200 |
| COLLAGEN IV | Abcam Ab-6586 | 1:100 |
| OTX2 | SCBT 30659 | 1:200 |
| FOXA2 | SCBT 6554 | 1:200 |
| PITX2 | Abcam Ab-55599 | 1:100 |
| EOMES | Abcam Ab-23345 | 1:100 |

### Chick immunofluorescence

Embryos were fixed in 4% paraformaldehyde in PBS for 1 hour at room temperature or overnight at 4°C. They were then washed 3 times with PBST (PBS+0.5% Triton X-100) for 1 hour each on nutator and blocked and permeabilized with 3% donkey serum, 1% bovine serum albumin in PBST for 2 hours, also at room temperature. Next, they were incubated overnight with anti-SOX2 antibody (R&D AF2018) diluted in blocking buffer at 4°C. The next day embryos were washed 3 times with PBST for 1 hour each on a nutator and then incubated with secondary donkey antibody Alexa-594, anti-human nuclear antigen (Novus Biologicals NBP2-34525AF647), and DAPI overnight. Embryos were washed times with PBST for 1 hour each and mounted in glass slides with fluoromount to image.

### Chick *in situ* hybridization

Chicken *SOX3* probe was kindly provided by F.M. Vieceli and the whole mount *in situ* hybridization was performed using previously described procedures^21^. Briefly, the embryos were fixed overnight in 4% paraformaldehyde in PBS 24-48 hours after the grafting. The embryos were then washed 3 times with PBS+0.1% Tween-20, and then dehydrated through a methanol series (25% methanol/PBS, 50% methanol/PBS, 75% methanol/PBS, 100% methanol), and rehydrated (100% methanol, 75% methanol/PBS, 50% methanol/PBS, 25% methanol/PBS PBS), 15 minutes each step at room temperature. Next, the embryos were incubated with Proteinase K 10μg/ml for 5 minutes, rinsed twice in PBS+0.1% Tween-20, incubated in 2mg/ml glycine in PBS+0.1% Tween-20, washed 2 times in PBS+0.1% Tween-20 for 5 minutes each and post-fix for 20 minutes in 4% paraformaldehyde +0.2% glutaraldehyde in PBS. The embryos were then hybridized at 70° C using antisense RNA chicken *SOX3, OTX2, HOXB1,* or *GBX2* probe labeled with digoxigenin-11-UTP. The probe was localized using AP-conjugated antibodies and the signal was developed with BM-Purple.

### Microscopy and Image Analysis

Images were acquired with either a Zeiss Axio Observer and a 20×/0.8 numerical aperture (NA) lens, or with a Leica SP8 inverted confocal microscope with a 40×/1.1-NA water-immersion objective. Image analysis and stitching was performed with ImageJ and custom Matlab routines. For the images used in Supplemental Video 1 and Supplemental Figure 8, these were also deconvolved with Autoquant software and analysed in Imaris. In these images the notochord-like feature was identified by a combination of manual and Ilastick classification based on DAPI morphology, and cells belonging to this structure were segmented and false-coloured with the assistance of custom Python 3D segmentation software written by J. Metzger.

### qPCR/RNAseq data

RNA was collected in Trizol at indicated time points from either mircopatterned colonies or from small un-patterned colonies. Total RNA was purified using the RNeasy mini kit (Qiagen). qPCR was performed as described previously^22^ and primer sequences are listed in the Supplemental Table 2. RNA-seq is from a previously published data set^22^, and all raw data are available from the GEO database, accession number GSE77057.

**Supplemental Table 2:**
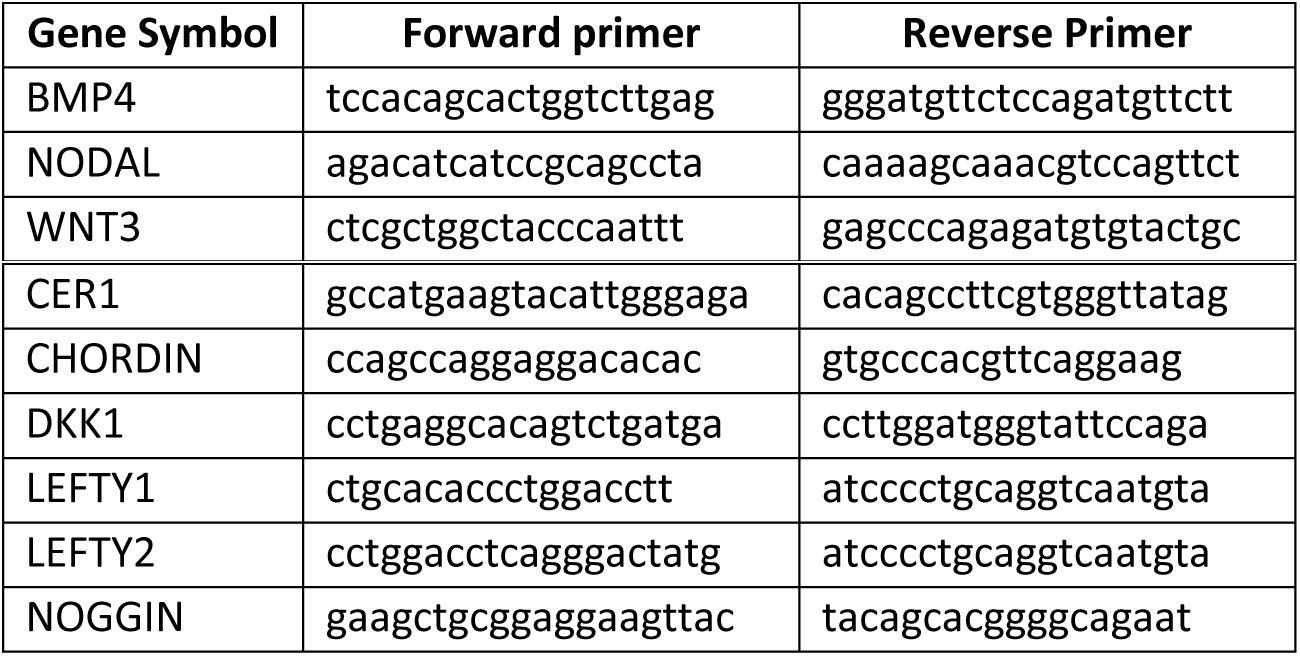
RT-QPCR Primer designs

### Transplantation of Human Organizer into Chick Host

Fertilized White Leghorn chicken eggs were incubated at 37-38°C and 50% of humidity and staged according to Hamburger and Hamilton^23^. Chick embryos were then removed from the egg and set up in Early-Chick culture^20^, with Pannett-Compton saline solution as final wash and residual liquid in the culture. Instead of growing on home-made micropatterned coverlsips, hESCs were grown on EMB Cytoo coverslips, as these had 500μm diameter micropatterns. All other culture details remained the same. Once grown to the indicated time with the indicated stimulation conditions, 500μm diameter colonies were peeled off whole with tungsten needles (Fine Science Tools). These colonies were washed twice with Pannett-Compton solution to remove culture growth factors and ligands. Colonies were then moved to chick embryos and grafted into the marginal zone between the area opaca and area pellucida, approximately 90° away from the site of primitive streak initiation. The grafted embryos were then returned to the incubator to develop and were imaged live one day after and were ultimately fixed between 24-48 hours post-graf. In all steps Pen/Strep was used to minimize the chance for bacterial contamination.

### Generation and validation of RUES2-GLR line

CRISPR/Cas9 technology was used to generate a single hESC line containing three independent fate reporters (SOX2-mCit, BRA-mCer and SOX17-tdTom). The already established and registered RUES2 hESC line (NIHhESC-09-0013) was used as the parental line. In order to achieve three independent targeting events in the same line, we approached each gene sequentially, since the efficiencies of recombination were not high enough for simultaneous targeting. First, for SOX17 targeting, we generated an homology donor plasmid (pSOX17-HomDon) containing:
i) a left homology arm that contain 1kb sequence right upstream of the SOX17 stop codon,
ii) a P2A-H2B-tdTomato cassette,
iii) a floxed Neomycin selection cassette (loxP-PGK-Neo-pA-loxP) and iv) a right homology arm containing a 1kb sequence immediately downstream of the SOX17 stop codon. Note that, since the H2B-tdTomato will be separated from the SOX17 gene by a self-cleaving P2A peptide, the fluorescent reporter will not be fused to SOX17, and therefore it will only be a reporter of the activation of the SOX17 expression, as the two proteins may have different half-lives. We initially tried a direct fusion of the SOX17 and tdTomato, but the fusion made the protein not properly localize to the nucleus, and therefore we decided to use a self-cleaving strategy. All DNA fragments were amplified from pre-existing plasmids or genomic DNA using Q5 polymerase with the primers listed in Supplementary Table 3, and joined together using standard DNA ligation protocols. An sgRNA recognizing a sequence near the stop codon of the SOX17 gene (Supplementary Table 4) was cloned into a Cas9-nickase expression vector (pX335 from the Zhang lab, Addgene plasmid #42335). This plasmid, together with the pSOX17-HomDon plasmid, were nucleofected into RUES2 cells using a Nucleofector II instrument and Cell Line Nucleofector Kit L (Lonza). Geneticin (a Neomycin analog) was added to the cultures 5 days after nucleofection, and kept in for 7 more days, to ensure selection of correctly targeted clones. Colonies derived from single geneticin-resistant cells were picked and expanded for screening. PCR amplification and Sanger-sequencing were used to identify correctly heterozygously targeted clones, with no unwanted mutations in the SOX17-sgRNA target site, both in the targeted and in the untargeted alleles. Positive clones were also validated for karyotyping (G-banding). The top 4 potential off-target sites of the sgRNA were PCR amplified and Sanger sequenced to ensure no unwanted mutations were present. The pluripotency status and absence of differentiation of the clones were validated through immunofluorescence staining. Once a perfectly validated SOX17-tdTom clone was identified, its youngest freezedown was thawed to undergo BRA gene targeting. Targeting of BRA followed a similar strategy as SOX17, but containing a Puromycin resistance cassette. Colonies derived from single puromycin-resistant cells were screened and validated as with the previous SOX17 targeting. After a fully validated double-targeted clone (SOX17-tdTom and BRA-mCer) was identified, it underwent sequential SOX2 targeting. Contrary to the SOX17 and BRA cases, for SOX2 targeting a direct fusion of SOX2 and mCitrine protein was possible without affecting its localization or function, and therefore the SOX2-mCitrine reporter constitutes a faithful reporter of both the “on” and “off” expression rates. The SOX2 homology donor consisted of: i) a 1kb left homology arm, ii) an mCitrine-T2A-Blasticidin cassette, and iii) a 1kb right homology arm. Colonies derived from single blasticidin-resistant cells were screened and validated as with the previous SOX17 and BRA targetings.

**Supplemental Table 3:**
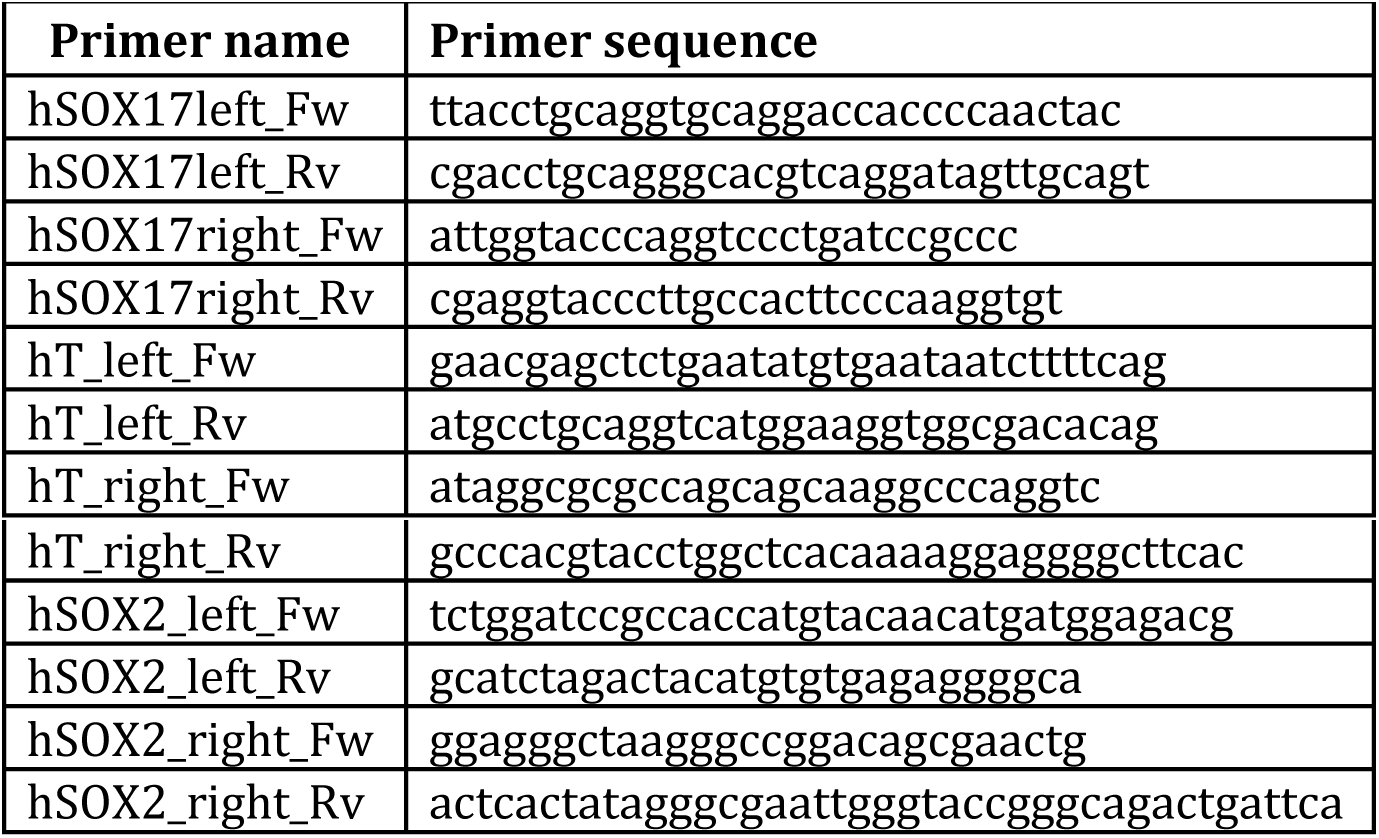
Primer sequences homology donor generation

**Supplemental Table 4:**
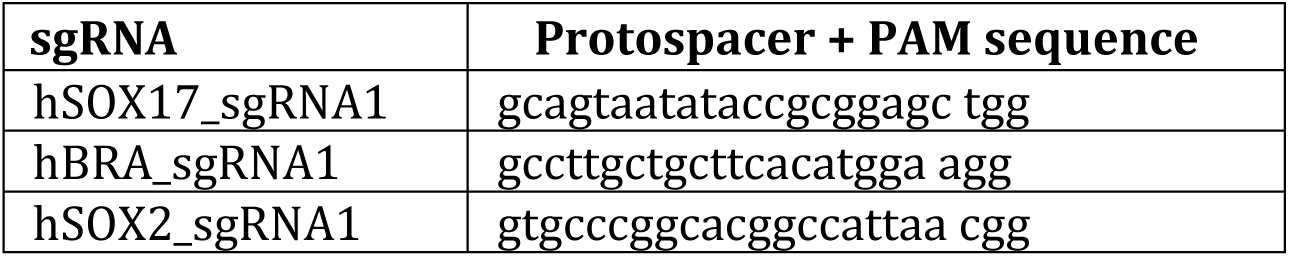
Sequences of the sgRNAs used for the generation of RUES2-GLR.

### RUES2-GLR time-lapse imaging

RUES2-GLR cells were dissociated to single cells from growth plates with StemPro Accutase (Life Technologies), washed, and then re-suspended in MEF-CM with 10μM ROCK-inhibitor Y-27632 (Abcam). CYTOO micropatterned chips were placed in 35mm tissue culture plastic dishes, and 8×10^5^ cells in 2mL of media were added to each coverslip. After 1 hour ROCK-inhibitor was removed and replaced with standard MEF-CM, supplemented with Pen/Strep (Life Technologies), and incubated overnight. The following morning the micropatterned coverslip was carefully removed from the dish and placed in a coverslip holder (CYTOOchambers from CYTOO), to which 1mL of MEF-CM+Pen/Strep+50ng/mL BMP4 was added to induce differentiation. Immediately after media addition, the holder was transferred to a spinning-disk confocal microscope (CellVoyager CV1000, Yokogawa), in which fluorescent images were acquired every 30 minutes for 2 days. Multichannel time-lapse movies were generated from the raw images using ImageJ analysis software.

## SUPPLEMENTAL FIGURE LEGENDS

**Supplemental Figure 1.**
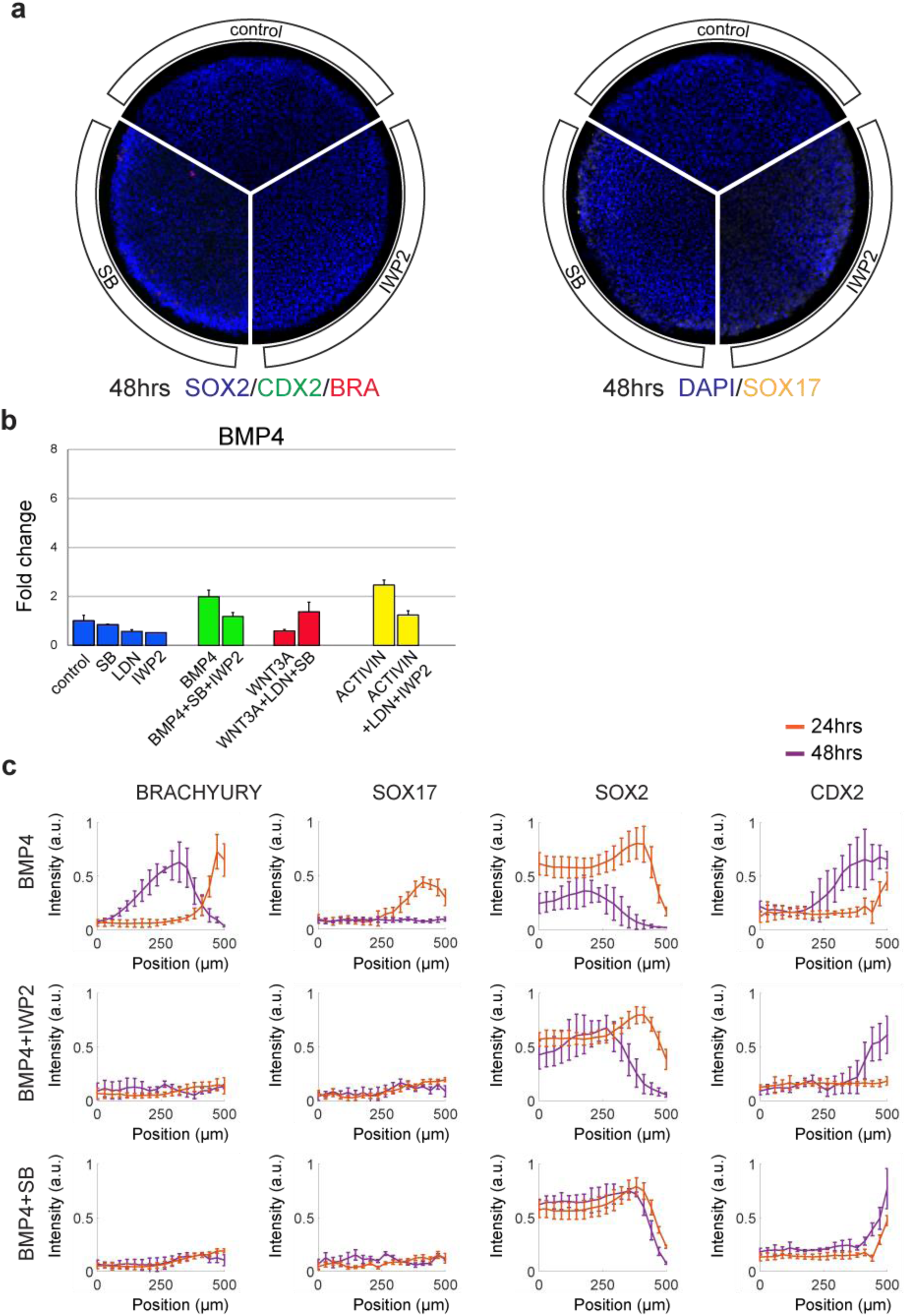
Controls for investigating hESC PS initiation hierarchy. (a) Micropatterned colonies stimulated with IWP2, SB, or blank media, fixed and stained for germ layer molecular markers after 48hrs. (b) qPCR for BMP4 of unpatterened small colonies stimulated for 4hrs conditions arraigned on the x-axis. As consistent with model hierarchy, there is no significant induction of BMP4 by ACTIVIN, WNT3A, or itself. (c) Quantification of Figure 1D. In this and in all other analysis unless stated otherwise, nuclei were segmented using DAPI and intensity of immunofluorescence signal for each marker was normalized to the DAPI intensity. Single cell expression data was binned radially and averaged. The final radial profile represents the average of *n*=25 colonies.

**Supplemental Figure 2.**
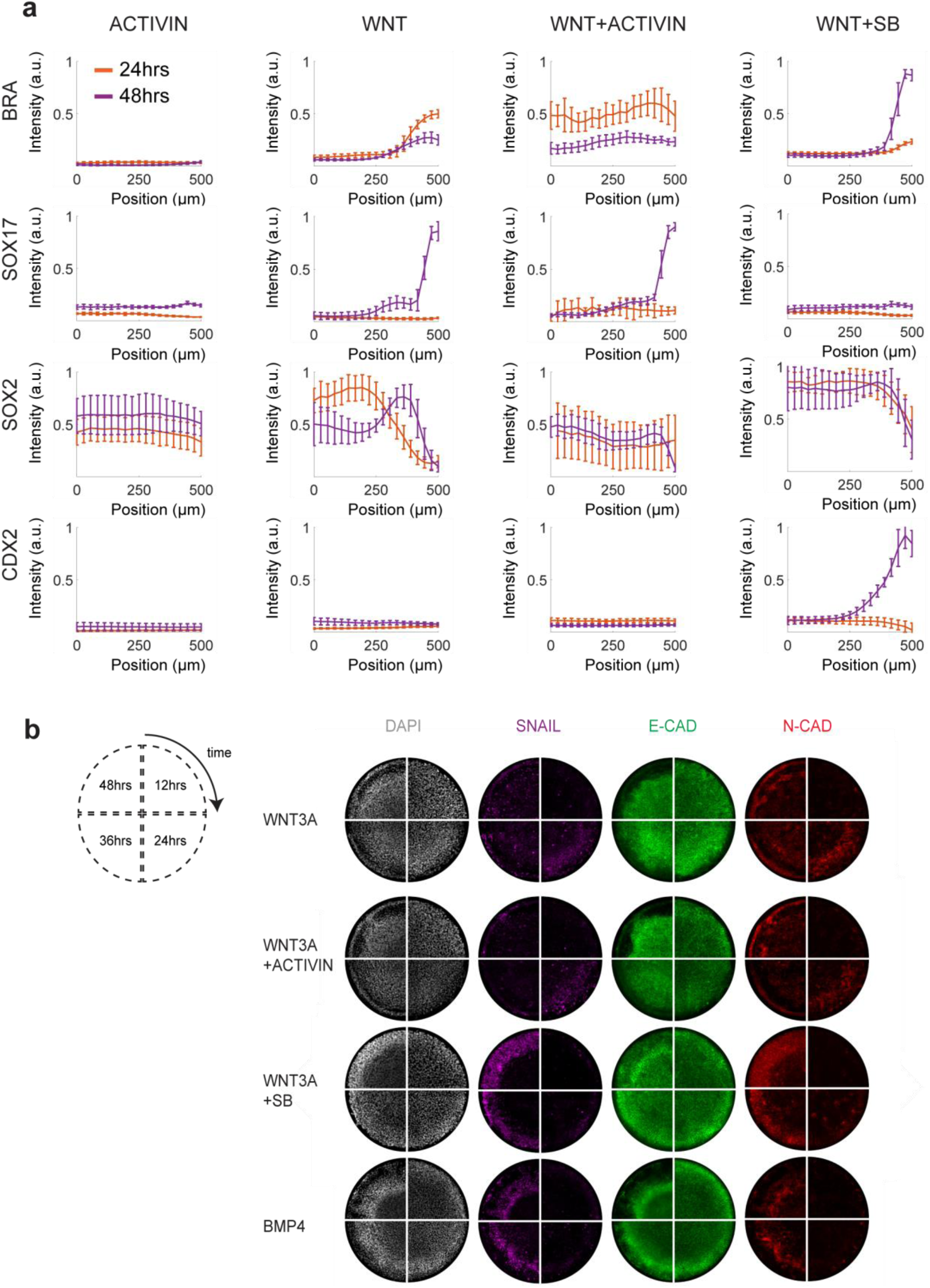
PS germ layer quantification and EMT timing. (a) Quantification of Fig. 2A. (b) Micropatterned colonies stimulated with BMP4, WNT3A, WNT3A+SB, or WNT3A+ACTIVIN and fixed and stained for primitive streak molecular markers SNAIL, E-CAD, and N-CAD after 12, 24, 36, or 48hrs. Note that WNT3A and WNT3A+ACTIVIN stimulated colonies turn on EMT markers faster than BMP4 or WNT3A+SB stimulated colonies, and have mostly downregulated SNAIL by 48hrs.

**Supplemental Figure 3.**
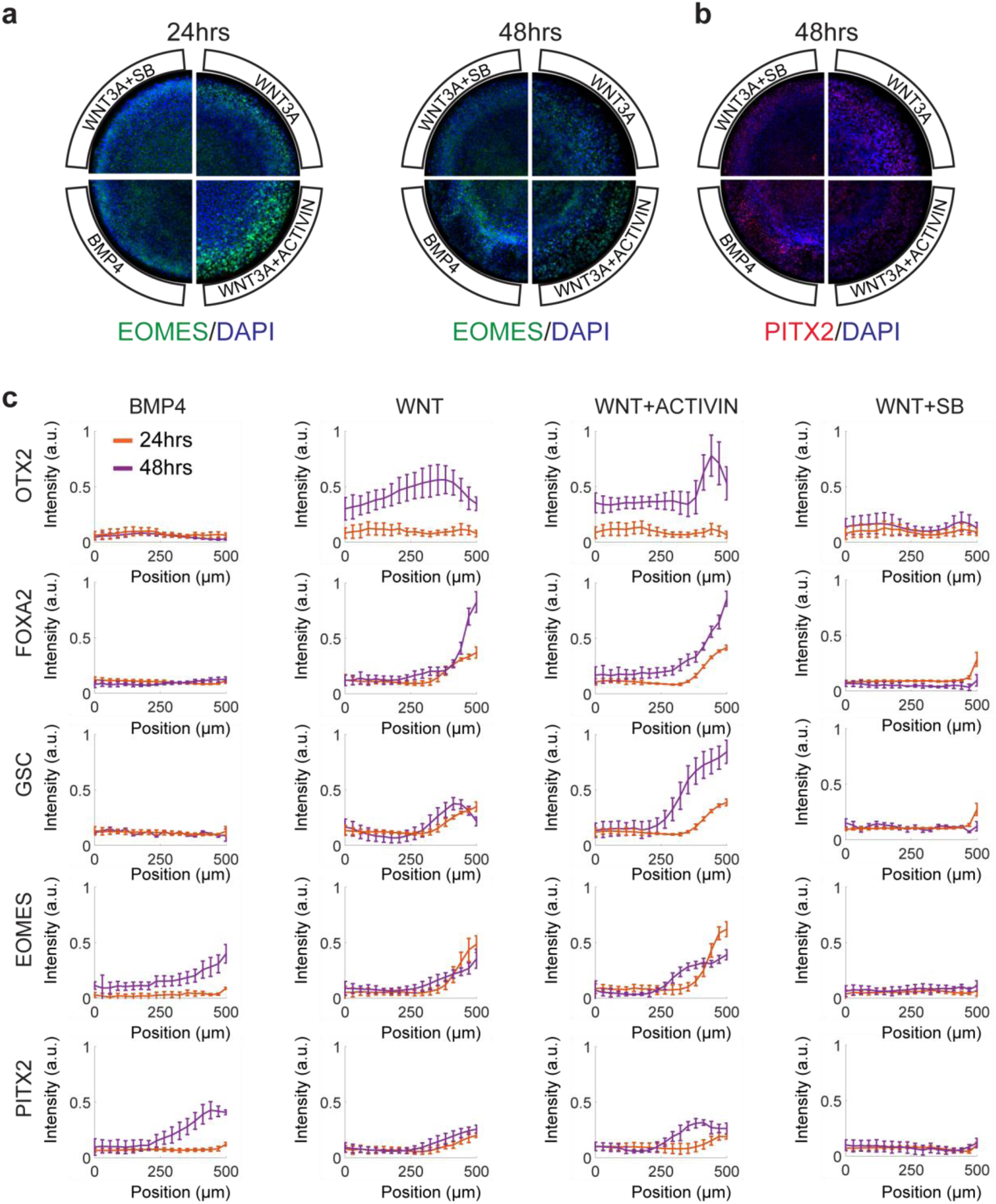
Further micropattern fate characterization. (a-b) Micropatterned colonies stimulated with BMP4, WNT3A, WNT3A+SB, or WNT3A+ACTIVIN and fixed and stained for EOMES at 24 and 48 hours (a) or PITX2 at 48 hours (b). EOMES is highest in WNT3A and WNT3A+ACTIVIN treated conditions and is also dynamic, with its highest expression at 24 hours, coinciding with the onset of PS markers (Supp. Fig. 2B). PITX2 is not highly expressed in any condition. (c) Quantification of Fig. 2J-K and Supp. Fig. 3A-B.

**Supplemental Figure 4.**
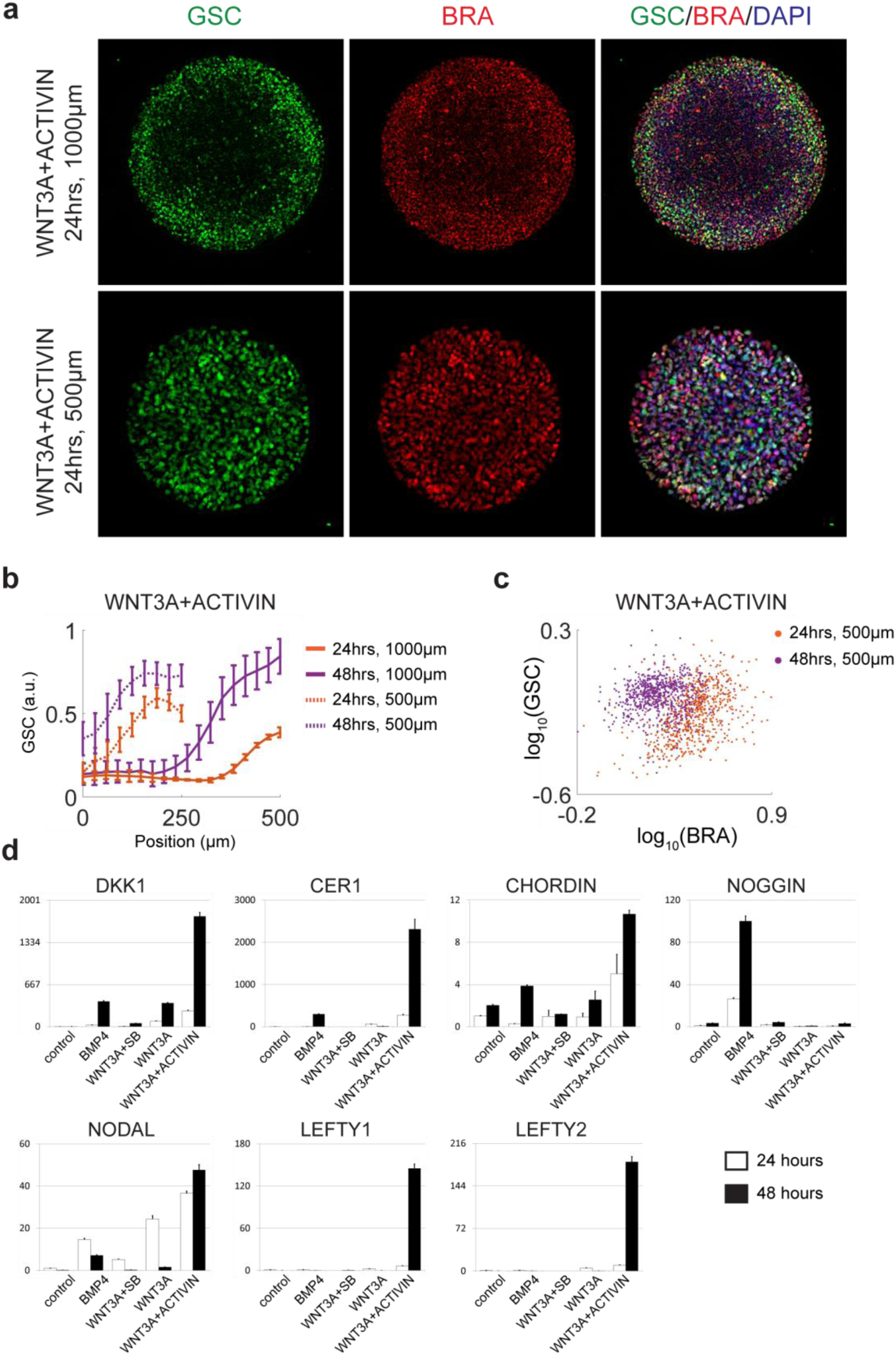
Further organizer characterization. (a) 1000μm and 500μm diameter micropatterned colonies stimulated with WNT3A+ACTIVIN and fixed and stained for GSC and BRA at 24 hours. Note that as observed by Warmflash et al.,^1^ for BMP induction, shrinking the colony size results in removal of center micropattern fate region, thus resulting here in a higher proportion of GSC expressing cells. (b) Quantification of (a). (c) Scatterplot of single-cell expression of GSC vs BRA. Note that at 24 hours most cells co-express BRA and GSC, but that by 48 hours GSC expression is increased and BRA expression is decreased. Because of this we grafted micropatterns at 24 hours as well as at 48 hours post-stimulation, reasoning that earlier coexpression of BRA and GSC would result in greater graft contribution to axial mesoderm structures. (d) qPCRs of additional organizer markers, taken from RNA collected from 500μm diameter micropatterns stimulated with either BMP4, WNT3A, WNT3A+SB, or WNT3A+ACTIVIN for 24 or 48 hrs. With the exception of NOGGIN, the characteristic organizer secreted inhibitors DKK1, CER1, CHORDIN, LEFTY1, and LEFTY2, are all most highly expressed in WNT3A+ACTIVIN conditions. The high NOGGIN induction by BMP4 in hESCs has been noted before^22^, and may represent a human-mouse species difference. NODAL, which in mouse is restricted to the organizer later in gastrulation, is also most highly expressed in WNT3A+ACTIVIN conditions.

**Supplemental Figure 5.**
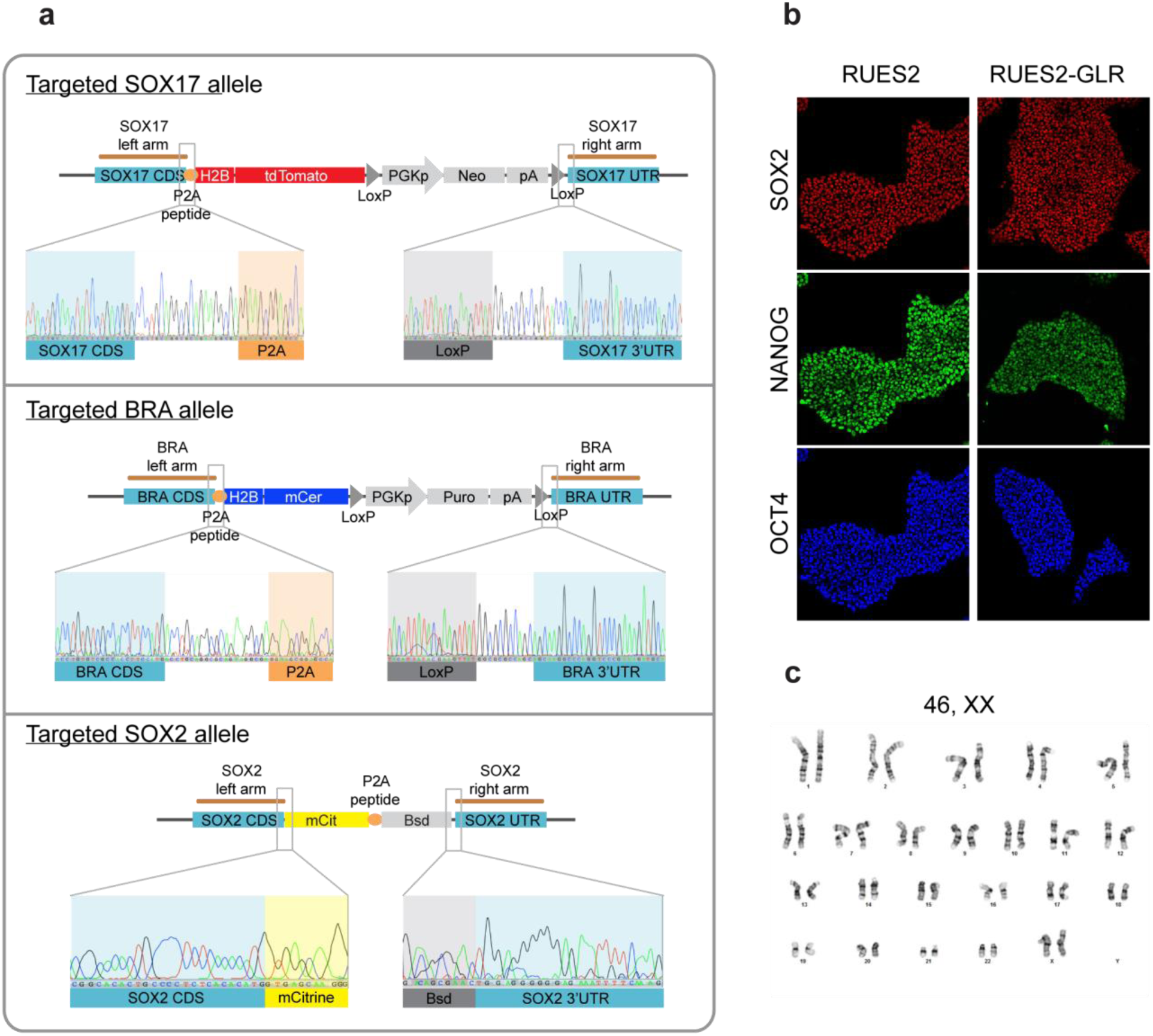
Generation and validation of RUES2-GLR cell line. (a) Sequencing of the targeted alleles of SOX17, BRA and SOX2 genes. No indels were detected. (b) The RUES2-GLR line maintain pluripotency normally, as assessed by the staining of typical pluripotency markers (OCT4, NANOG and SOX2). Scale bar is 100μm. (c) The RUES2-GLR line was karyotypically normal.

**Supplemental Figure 6.**
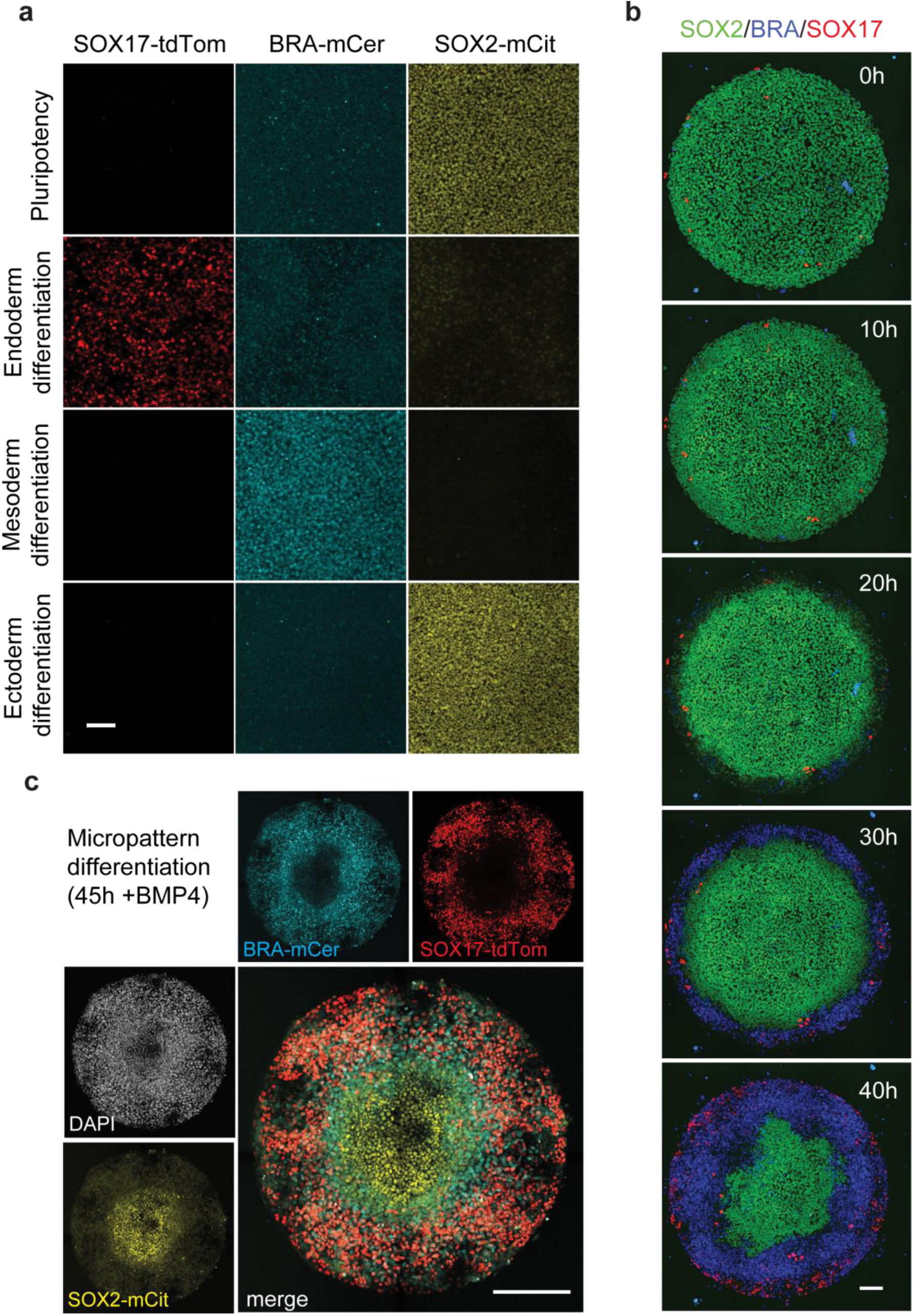
Functional validation of RUES2-GLR cell line. (a) Specificity of the germ layer reporters. When induced to differentiate to individual germ fates, only the specific reporter was turned on. SOX2-mCitrine reporter was expressed during pluripotency and 3 days after neural (ectoderm) differentiation, BRA-mCerulean was turned on after 3 days of mesodermal differentiation and SOX17-tdTomato reporter was active after 3 days of endodermal differentiation. Scale bar is 100μm. (b) Snapshots of a time-lapse imaging of the RUES2-GLR line after 50ng/mL BMP4 in micropatterns, showing how differentiation starts from the edges and extends inwards. Scale bar is 100μm. (c) RUES2-GLR line reproducibly generated the typical self-organized concentric rings of germ layers when induced to differentiate with a step presentation of 50ng/mL BMP4 in micropatterns. Scale bar is 200μm.

**Supplemental Figure 7.**
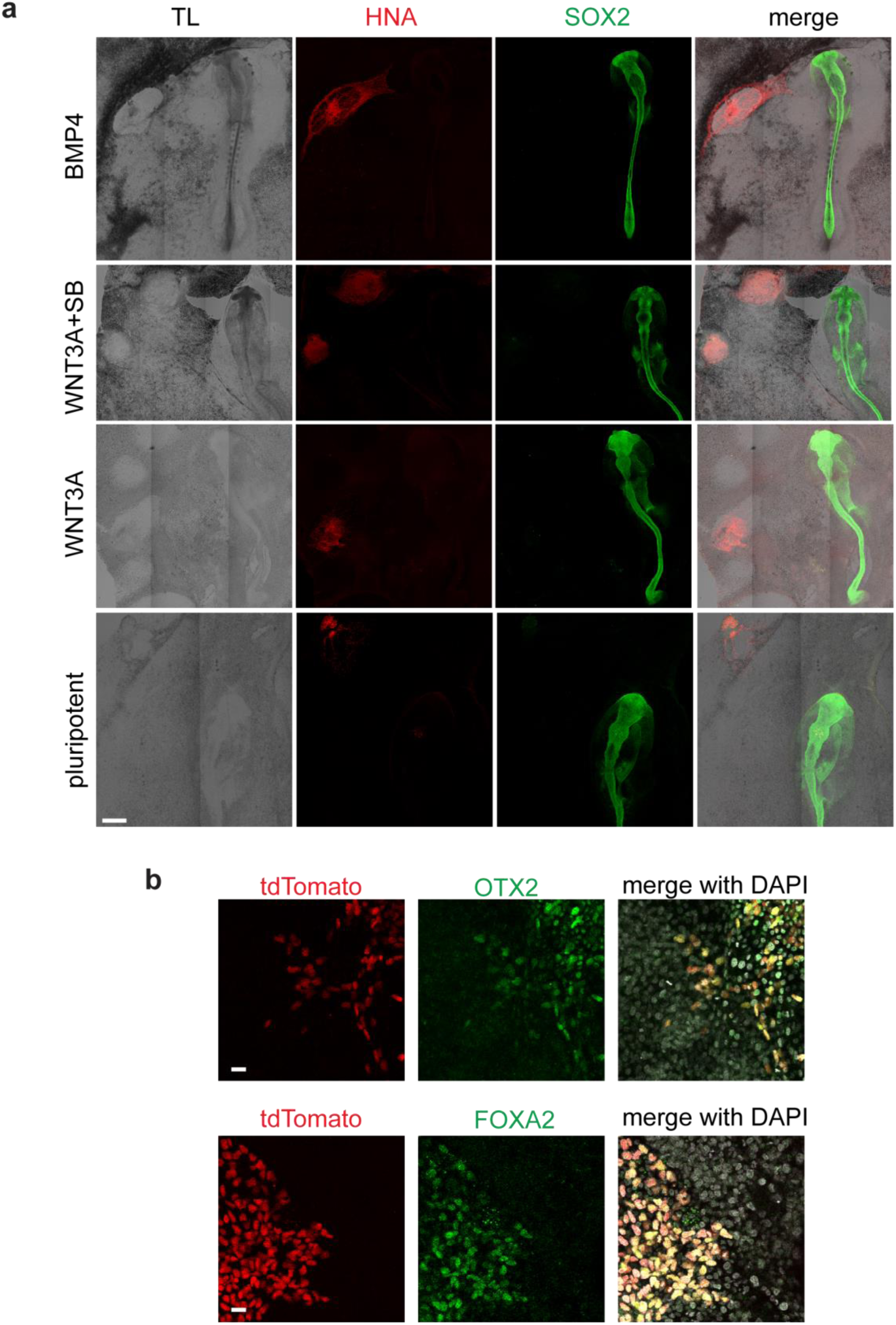
Control chick grafts. (a) Representative grafts for control conditions. With the exception of the BMP4 control condition, grafted hESC colonies were static, with the colonies either growing or dying in place. With BMP4, often the colonies were elongated, possibly due to hESC migration. In all control conditions, however, there was never induction of SOX2 in the host cells. Note that in the case of the WNT3A+SB graft shown, two colonies were grafted into two different locations. Scale bar is 500μm. (b) Confocal cross-sections showing co-expression of SOX17 (tdTomato) and FOXA2 or OTX2 in human cells that contribute to the secondary axes induced by a 24hr WNT3A+ACTIVIN stimulated hESC micropattern. Scale bar is 20μm.

**Supplemental Figure 8.**
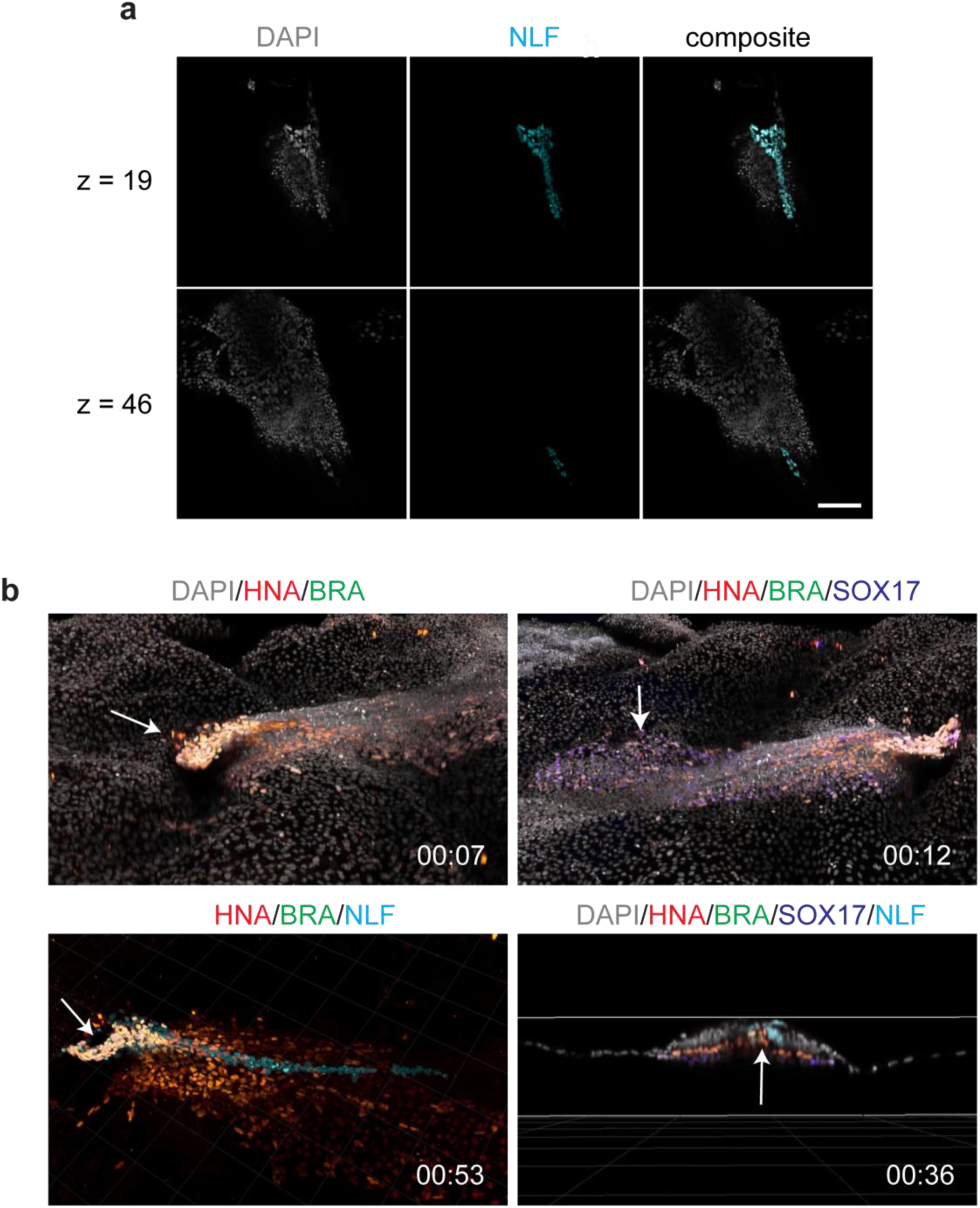
Stills of Supplemental Video 1. (a) Examples of classifying the notochord-like feature (NLF) based on morphology. For z=+19μm, one can discern the NLF as a tighter and brighter rod of cells running north-south that is also distinct and somewhat separated from the surrounding chick epiblast. For z=+46μm, one sees that paired elongated cells stick out ahead of the other cells in a continuation of the originally identified NLF. Other cells belonging to the NLF between z=+46μm and z=+19μm are obscured at these slices or out of focus, but can be easily identified slice-by-slice at the other z positions. Scale bar is 100μm. (b) Snapshots of Supplemental Video 1. Clockwise from top-left: yellow shows colocalization of BRA (green) and human (red) cells; purple shows co-localization of Sox17:tdTomato (blue) with human (red) cells; cross-section shows that chick and human cells arrange themselves into germ layers properly, and that they flank the central notochord-like feature indicated by the arrow (cyan); a proportion of human mesoderm cells contribute to part of the notochord-like structure, while the cyan-coloured cells without HNA (red) shows that the remainder of the NLF is composed of host cells.

**Supplemental Video 1**

Fly-through animation of confocal z-stack acquired from the embryo shown in Figure 3L-R. Stack was first deconovled with Autoquant software, and then manipulated in Imaris. For classification methodology of the notochord-like feature (NLF) please see Methods and Supplemental Figure 8.

